# *rescomp*: An R package for defining, simulating and visualizing ODE models of consumer-resource interactions

**DOI:** 10.1101/2022.01.11.475574

**Authors:** Andrew D. Letten

## Abstract

1. Mechanistic models of resource competition underpin numerous foundational concepts and theories in ecology, and continue to be employed widely to address diverse research questions. Nevertheless, current software tools present a comparatively steep barrier to entry.
2. I introduce the R package *rescomp* to support the specification, simulation and visualisaton of a broad spectrum of consumer-resource interactions. *rescomp* is compatible with diverse model specifications, including an unlimited number of consumers and resources, different consumer functional responses (type I, II and III), different resource types (essential or substitutable) and supply dynamics (chemostats, logistic and/or pulsed), delayed consumer introductions, time dependent growth and consumption parameters, and instantaneous changes to consumer and/or resource densities.
3. Several examples on implementing *rescomp* are provided. In addition, a wide variety of additional examples can be found in the package vignettes, including using *rescomp* to reproduce the results of several well known studies from the literature.
4. *rescomp* provides users with an accessible tool to reproduce classic models in ecology, to specify models resembling a wide range of experimental designs, and to explore diverse novel model formulations.

## Introduction

There is a rich tradition in ecology of using models of resource competition described by systems of ordinary differential equations (ODEs) to explore species interactions and their community level consequences (Lafferty *et al*. 2015). Consumer-resource models have been the basis for numerous classical concepts and theories, including Tilman’s resource-ratio theory (Tilman 1982), apparent competition (Holt 1977), the gleaner-opportunist trade-off (Grover 1997), the paradox of enrichment (Rosenzweig 1971), diversity maintenance via chaos (Huisman & Weissing 1999), and the hydra effect (Abrams 2009), to name just a few. They continue to be employed widely to address a growing diversity of questions, including research spanning the gut microbiome (Guittar *et al*. 2021), antibiotic resistance (Levin & Udekwu 2010; Nev *et al*. 2020; Letten *et al*. 2021), cancer biology (Huntly *et al*. 2021), and collective behaviour (Dalziel *et al*. 2021). In addition, they are routinely drawn upon in textbooks and undergraduate teaching material in ecology and evolution (Morin 2011; Otto & Day 2011).

Despite the central role consumer-resource models play both in theory development and in the teaching of foundational concepts in ecology, a considerable level of technical knowledge (mathematical and programming) is required for their specification and simulation using current software tools. This is particularly true for models incorporating non-autonomous behaviour such as time-dependent parameters and discontinuities in state variables. The R package *deSolve* provides a powerful and highly flexible interface for simulating systems of ODEs, but requires the user to explicitly specify the model equations and any auxiliary functions (e.g., those regulating non-autonomous behaviour). This can present a relatively steep barrier to entry to practitioners unfamiliar with the defining of ODE models and/or writing custom functions in R.

Here, I introduce *rescomp* (https://andrewletten.github.io/rescomp/), an R package that facilitates the specification, simulation and visualisaton of a broad spectrum of consumer-resource competition models. At its core, *rescomp* provides a resource competition focused interface to *deSolve*, alongside flexible functions for visualizing model properties and dynamics. A diverse range of model properties and behaviour are supported by *rescomp*, including an unlimited number of consumers and resources, different consumer functional responses, different resource types (e.g., essential or substitutable) and supply dynamics, delayed consumer introductions, time dependent growth and consumption parameters, and instantaneous changes to consumer and/or resource densities (e.g., in serial batch transfer). This provides users with a convenient tool to reproduce numerous classic models in ecology; to specify models resembling a wide range of experimental designs; and to explore diverse novel model formulations.

### A quickstart guide to *rescomp*

The primary user function in *rescomp* is `spec_rescomp`, which facilitates: i) the definition and parameterisation of a consumer-resource model, and ii) the specification of simulation parameters.

The default output from `spec_rescomp` is an S3 object of class `rescomp` defining a model for a single type I consumer (linear functional response) and a single continuously supplied resource (e.g., in a chemostat). By default, calls to `spec-rescomp` print a descriptive summary of the model and simulation specifications (to inhibit printing of the summary use `verbose = FALSE`).

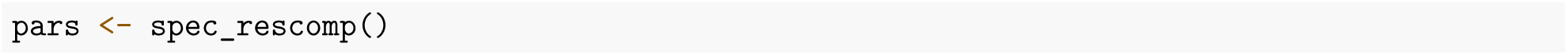

~~~
## Model properties
## * 1 consumer(s) and 1 resource(s)
## * Consumers have type 1 functional responses
## * Resource supply is continuous (e.g. chemostat)
## * Mortality is continuous
##
## Simulation properties
## * Simulation time: 1000 time steps
## * Init state: consumer(s) = [10], resource(s) = [1]
~~~

The remaining key functions in *rescomp* operate downstream from `spec_rescomp. plot_funcresp` plots the functional response(s) for easy visualistion prior to running a simulation (Fig 1a). The model can then be simulated via `sim_rescomp`, which is effectively a wrapper for `deSolve::ode` with convenient defaults. The resultant output dynamics are efficiently visualised with `plot_rescomp` (Fig 1b).

**Figure 1:**
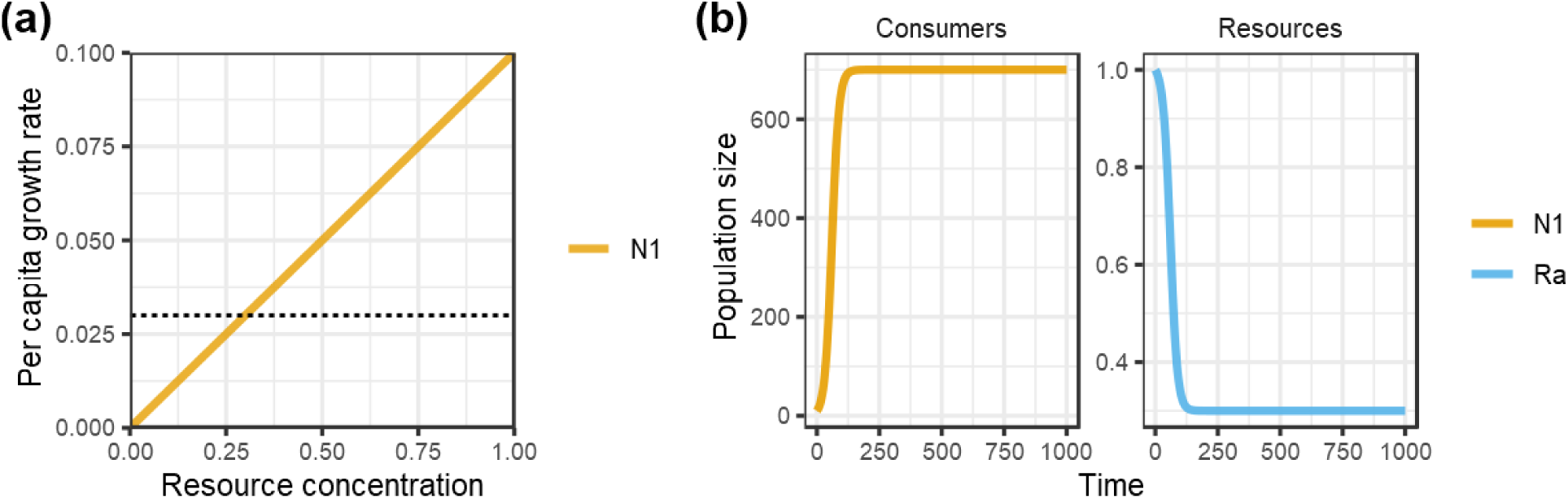
Default output from *rescomp* plotting functions. **a)** The function `plot_funcresp` plots the functional responses of the consumers for a specified model returned by `spec_rescomp`, in this instance a single consumer with a linear (type I) functional response. **b)** The function `plot_rescomp` plots the corresponding consumer and resource dynamics outputted by `sim_rescomp.`

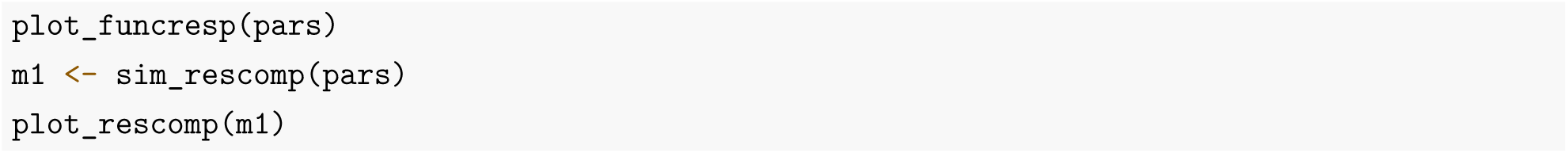

Note, the core `rescomp` functions are also compatible with base R pipes, such that `spec_rescomp() |> sim_rescomp() |> plot_rescomp()` will output Figure 1b.

The main utility of *rescomp* comes with specifying more elaborate models and simulation dynamics, which can be tailored to a broad range of study systems and research questions. Before exploring these options in more depth, the following section provides a brief summary of the mathematical structure of consumer-resource models in *rescomp*.

### Model formulation

All *rescomp* models take the general form,

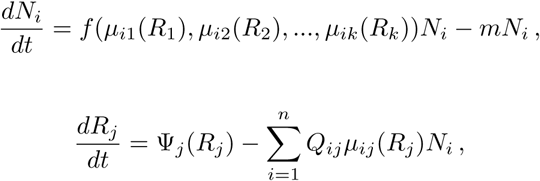

where *N*_*i*_ is the population density of consumer *i, R*_*j*_ is the density/concentration of resource *j, μ*_*ij*_ is the per capita consumer functional response of consumer *i* on resource *j, m* is the per capita mortality rate (constant or pulsed), Ψ_*j*_(*R*_*j*_) is the resource supply function, and *Q*_*ij*_ is the resource quota of consumer *i* on resource *j* (the amount of resource per unit consumer). From this general form, different model formulations are distinguished by: i) the number of consumers/resources; ii) the form of the consumer functional response; iii) the mode of resource supply; iv) the type of resource; and v) any non-autonomous behaviour including time dependent model parameters and/or instantaneous changes in consumer/resource density (e.g., in batch transfer). In the case of multiple resources, each resource is either treated as essential to consumer growth following Leibig’s law of the minimum, in which case, *f* (*μ*_*i*1_(*R*_1_), *μ*_*i*2_(*R*_2_), *…, μ*_*ik*_(*R*_*k*_)) = min(*μ*_*i*1_(*R*_1_), *μ*_*i*2_(*R*_2_), *…, μ*_*ik*_(*R*_*k*_)); or substitutable such that: *f* (*μ*_*i*1_(*R*_1_), *μ*_*i*2_(*R*_2_), *…, μ*_*ik*_(*R*_*k*_)) = *f* (*μ*_*i*1_(*R*_1_) + *μ*_*i*2_(*R*_2_) + … + *μ*_*ik*_(*R*_*k*_)).

#### Consumer equations

The consumer growth function can take one of three forms: i) linear (type I), *μ*_*ij*_(*R*_*j*_) = *a*_*ij*_*R*_*j*_, where *a*_*ij*_ is a resource consumption constant; ii) nonlinear type II (aka Monod), 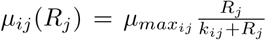, where *μ*_*maxij*_ is the maximum growth rate and *k*_*ij*_ is the half saturation constant for consumer *i* on resource *j*; or iii) nonlinear type III (sigmoidal), 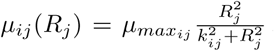. An alternative parameterisation to all three functional responses is to incorporate a consumption efficiency term, *c*_*ij*_. The type I model becomes *μ*_*ij*_(*R*_*j*_) = *a*_*ij*_*c*_*ij*_*R*_*j*_, and for type II and III, *a*_*ij*_*c*_*ij*_ is substituted for *μ*_*max*_. Note this functional form is mutually exclusive with the specification of resources quotas in the resource equations, i.e., the *Q*_*ij*_s are dropped from the resource equations.

#### Resource equations

The resource supply function, Ψ_*j*_(*R*_*j*_), can also take several forms. Specifically, either the resources are biological and grow logistically, 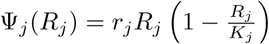, where *r*_*j*_ is the resource intrinsic rate of increase and *K*_*j*_ is the resource carrying capacity; or the resources are supplied to the system at a fixed concentration and rate (as in a chemostat), Ψ_*j*_(*R*_*j*_) = *d*(*S*_*j*_ − *R*_*j*_), where *d* represents the flux of resources into and out of the system; or the resources are pulsed intermittently.

### Further options, arguments and examples

As a simple extension of the default arguments provided by `spec_rescomp`, the following code defines a model for two consumers consuming two substitutable resources, with the corresponding plots returned by `plot_funcresp` and `plot_rescomp` shown in Figure 2 (a + b).

**Figure 2:**
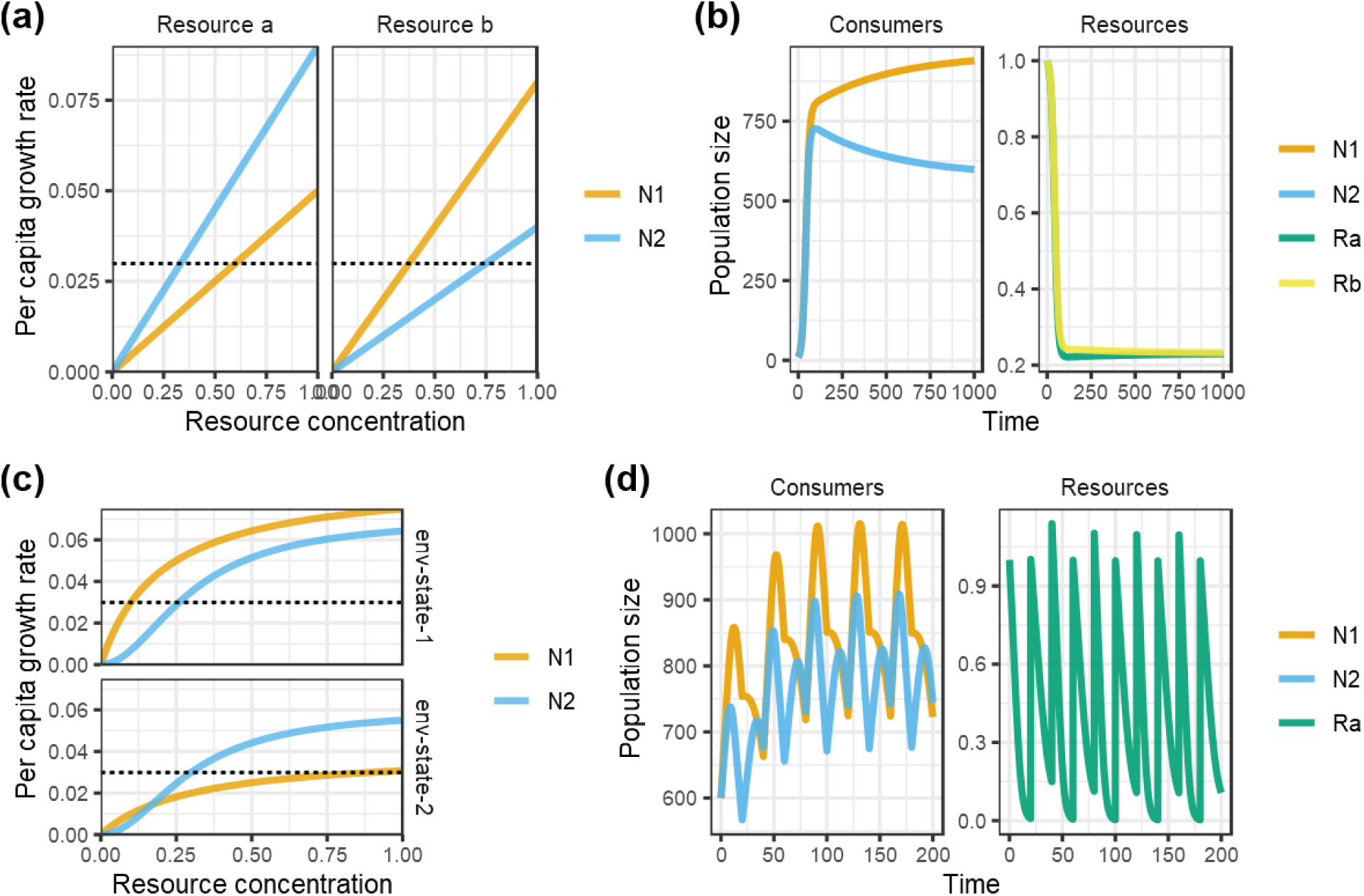
Visualizing functional responses and dynamics of consumer resource models with multiple consumers and/or resources: **a**) and **b)** two consumers and two substituable resources; **c)** and **d)** two consumers with time-dependent consumption parameters and a single pulsed resource. **a)** When multiple resources are specified in `spec_rescomp, plot_funcresp` will facet by resource. **b**)The corresponding consumer and resource dynamics outputted by `sim_rescomp`. **c)** When time dependent parameters are specified in `spec_rescomp, plot_funcresp` will facet by alternative parameter states. **d)** The corresponding consumer and resource dynamics outputted by `sim_rescomp`, where the instantaneous resource pulses *and* the switch between environmental states are specified to coincide every 20 time-steps.

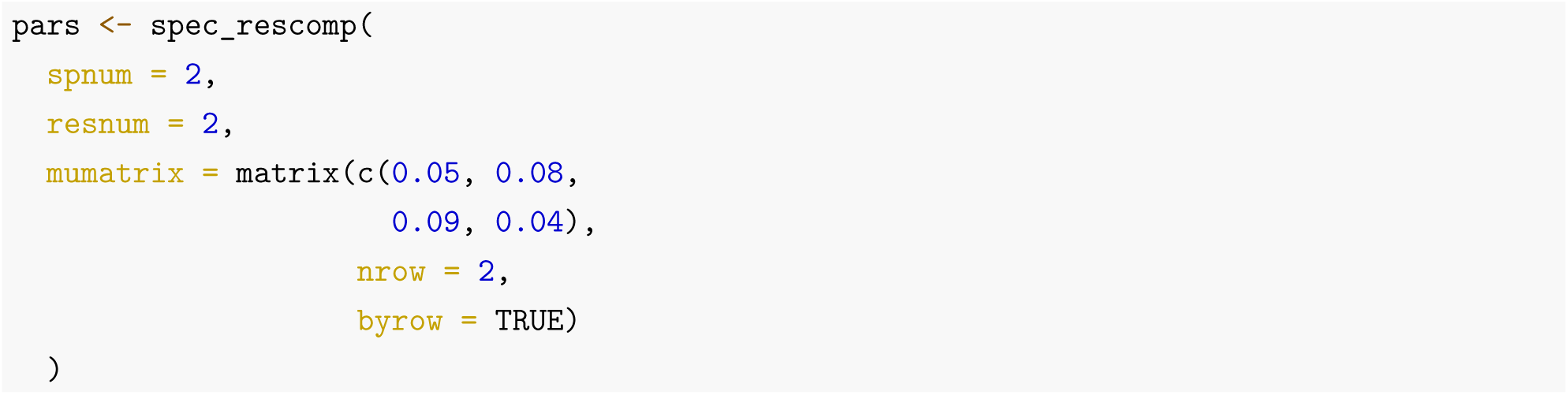

~~~
## Model properties
## * 2 consumer(s) and 2 resource(s)
## * Consumers have type 1 functional responses
## * Resources are substitutable
## * Resource supply is continuous (e.g. chemostat)
## * Mortality is continuous
##
## Simulation properties
## * Simulation time: 1000 time steps
## * Init state: consumer(s) = [10, 10], resource(s) = [1, 1]
~~~

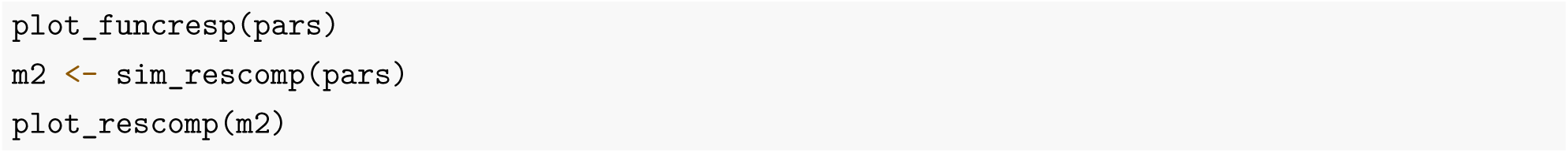

Beyond specifying the number of consumers and resources, consumer functional responses, and the type of resource, *rescomp* users can also choose to incorporate more complex model attributes and behaviour, such as time-dependent parameters, instantaneous changes in state variable abundance/density and/or delayed introductions (a brief summary of all arguments available in `spec_rescomp` is provided in Table 1). For example, the following code defines a model for two consumers (type II and type III), with time dependent growth parameters, competing for a single pulsed resource. Time dependent growth parameters facilitate the modeling of non-constant environmental conditions, such as a variable experimental treatment or seasonal dynamics. Resource pulsing allows the user to specify the exact time at which resources are supplied to the system, capturing, for example the resupply of resources in experimental batch culture or the intermittent pulsing of resources in the mammalian gut. The consumer functional responses and output dynamics for this model, as generated by `plot_funcresp` and `plot_rescomp` respectively, are shown in Figure 2 (c&d).

**Table 1:** Summary of function arguments available in `spec_rescomp.`

| Argument | Description |
| --- | --- |
| `spnum` | Number of consumers |
| `resnum` | Number of resources |
| `mumatrix` | Maximum growth rates / resource consumption constants |
| `kmatrix` | Half saturation constants |
| `qmatrix` | Resource quotas |
| `effmatrix` | Resource conversion efficiencies |
| `funcresp` | Functional response (type I, II or III) |
| `essential` | Essential or substitutable resources |
| `chemo` | Chemostat or logistic resource supply |
| `mort` | Mortality rates |
| `resspeed` | Resource supply / growth rate |
| `resconc` | Resource concentration / carrying capacity |
| `respulse` | Resource pulse size |
| `mortpulse` | Pulsed mortality fraction |
| `pulsefreq` | Resource / mortality pulse frequency |
| `batchtrans` | Implement serial transfer in batch culture |
| `timepars` | Implement time-dependent growth / consumption parameters |
| `totaltime` | Total simulation length |
| `timeparfreq` | Frequency of switching between alternative time-dependent parameters |
| `tpinterp` | Method for interpolating time-dependent parameters |
| `cinit` | Consumer starting densities |
| `rinit` | Resource starting concentrations |
| `introseq` | Consumer introduction times |
| `verbose` | Print model summary |

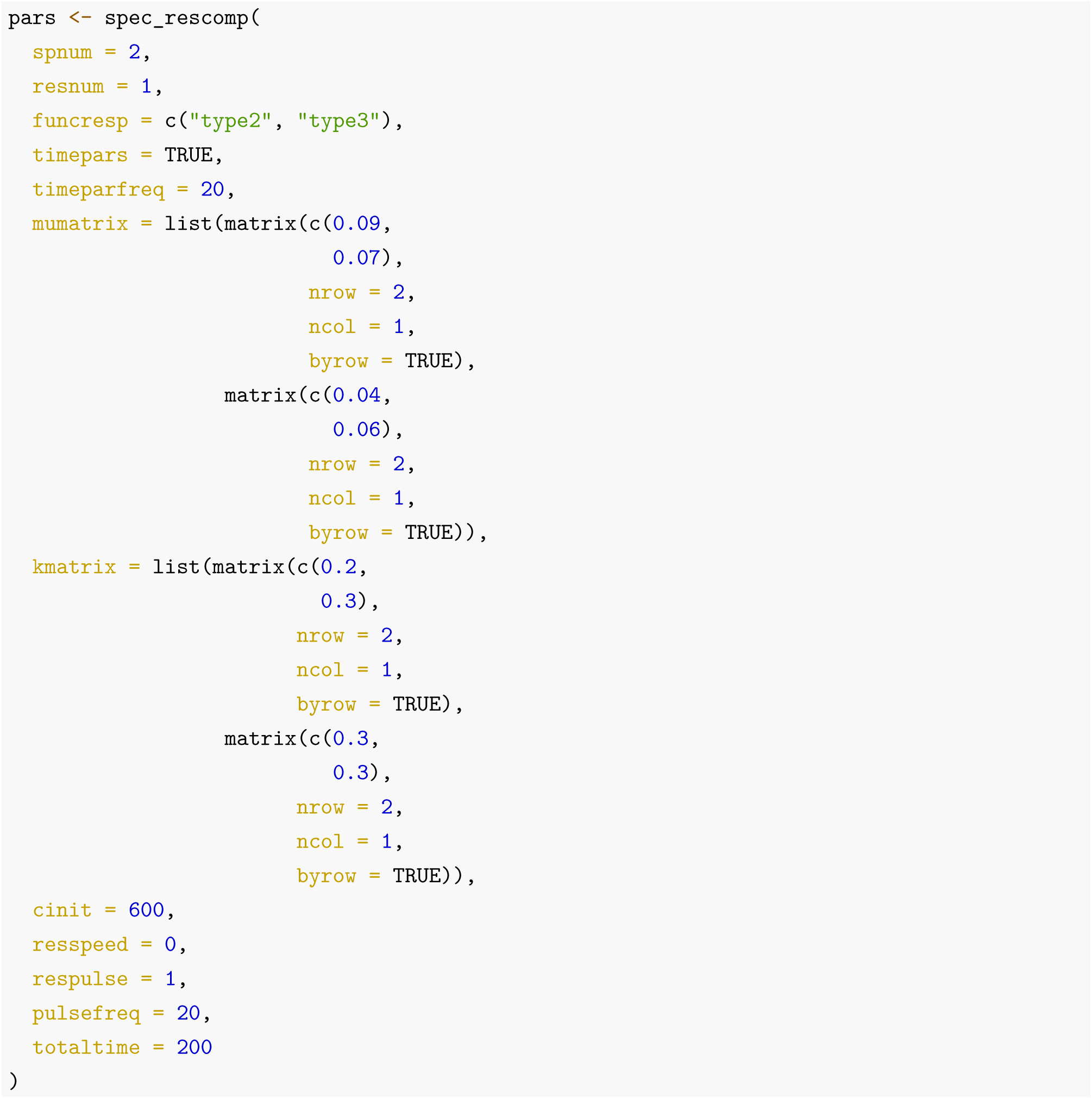

~~~
## Model properties
## * 2 consumer(s) and 1 resource(s)
## * Consumers have type 2 or type 3 functional responses
## * Resource supply is pulsed only
## * Mortality is continuous
## * Time dependent parameters with instantaneous switching every 20 timesteps
##
## Simulation properties
## * Simulation time: 200 time steps
## * Resources pulsing every 20 timesteps
## * Init state: consumer(s) = [600, 600], resource(s) = [1]
~~~

A wide range of other examples can be found in the package vignettes (https://andrewletten.github.io/rescomp/articles/rescomp.html, https://andrewletten.github.io/rescomp/articles/classic-results.html). The vignette *Reproducing studies in resource competition with rescomp* illustrates the use of *rescomp* to reproduce various classic results in resource competition from the literature, including Huisman & Weissing (1999), Grover (1990) and Armstrong & McGehee (1980). Examples taken from Huisman & Weissing (1999) include six species consuming three essential and continuously supplied resources (Fig 3a), and five species exhibiting chaotic dynamics on five essential resources (Fig 3b). Implementing the examples from Huisman & Weissing (1999) requires the use of the introseq argument in `spec_rescomp`, which enables delayed consumer introduction times. The example taken from Armstrong & McGehee (1980) illustrates the coexistence of a type I consumer and a type II consumer on a single logistically growing resource (Fig 3c&d). Note that Armstrong and McGehee parameterised their model using conversion efficiencies in the consumer equations rather than quotas in the resource equations. In `spec_rescomp` this is implemented via the specification of an `effmatrix` rather than a `qmatrix`.

**Figure 3:**
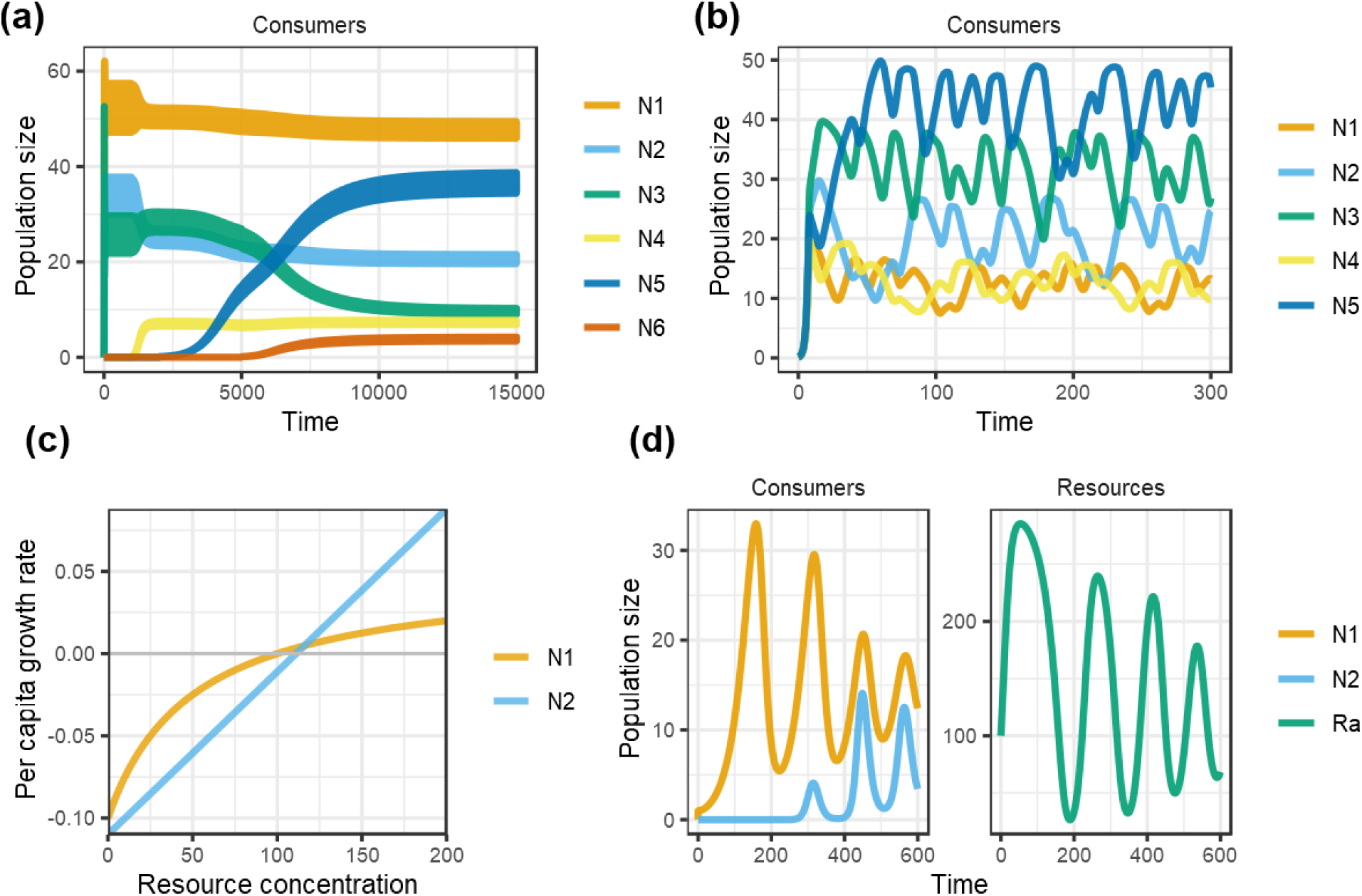
Figures from package vignette - *Reproducing studies in resource competition with rescomp*. **a)** Reproduces Fig 1c from Huisman and Weissing (1999). **b)** Corresponds to Fig 2a of Huisman and Weissing (1999). **c)**. Corresponds to Fig 2 from Armstrong and McGehee (1980). Because the two consumers were specified with slightly different density independent mortality rates, it is more informative to plot the functional responses using the `madj = TRUE` argument in `plot_funcresp`. **d)** Corresponds to Fig 1 from Armstrong and McGehee (1980). The paper states that the model was initiated with R = 400, but here it is started with R = 100 to provide as close as possible correspondence to the original figure.

### Conclusion

The goal of *rescomp* is to streamline the process of defining, simulating and visualizing ODE models of ecological consumer-resource interactions. *rescomp* is designed to support both theoretical investigations by facilitating the flexible exploration of novel model formulations and/or simulation dynamics; and empirical research by providing experimenters with an accessible tool for generating model-based hypotheses and for interpreting results. In addition, *rescomp* provides instructors with an interactive application to introduce students to basic models in resource competition as well as for reproducing classic studies from the literature.

